# Sparse interaction between oligodendrocyte precursor cells (NG2^+^ cells) and nodes of Ranvier in the central nervous system

**DOI:** 10.1101/185801

**Authors:** Lindsay M. De Biase, Michele L. Pucak, Shin H. Kang, Stephanie N. Rodriguez, Dwight E. Bergles

## Abstract

Regeneration of propagating action potentials at nodes of Ranvier allows nerve impulses to be conducted over long distances. Proper nodal function is believed to rely on intimate associations among axons, myelinating oligodendrocytes, and perinodal astrocytes. Studies in the optic nerve, corpus callosum, and spinal cord suggest that NG2^+^ cells are also key constituents of CNS nodes and that these glia may influence conduction efficacy and formation of axon collaterals. However, the prevalence of NG2^+^ cell processes at CNS nodes of Ranvier has not been rigorously quantified. Here we used a transgenic mouse expressing membrane-targeted EGFP to visualize the fine processes of NG2^+^ cells and to quantify the spatial relationship between NG2^+^ cells and nodes of Ranvier in four distinct CNS white matter tracts. NG2^+^ cell processes came within close spatial proximity to a small percentage of nodes of Ranvier and approximately half of these spatial interactions were estimated to occur by chance. The majority of NG2^+^ cell process tips were not found in close proximity to nodes and gray matter NG2^+^ cells in regions of low nodal density were as morphologically complex as their white matter counterparts, indicating that attraction to nodes does not critically influence the elaboration of NG2^+^ cell processes. Finally, there was no difference in nodal density between small regions devoid of NG2^+^ cell processes and those containing numerous NG2^+^ cells processes, demonstrating that the function of CNS nodes of Ranvier does not require ongoing interaction with NG2^+^ cells.

**Significance Statement:** Effective propagation of action potentials along neuronal axons is dependent upon periodic regeneration of depolarization at nodes of Ranvier. The position, structural integrity, and function of nodes of Ranvier is believed to be regulated, in part, by intimate physical interactions between nearby glial cells and nodes. Clarifying whether oligodendrocyte precursor cells are obligate members of this nodal support system is critical for defining whether these cells contribute to pathologies in which nodal structure is compromised.

## INTRODUCTION

Conduction velocity in central axons is exquisitely regulated by axon diameter, nodal size, nodal ion channel composition, and the length of internodes, or the region of compact myelin between nodes of Ranvier (NOR). Indeed, these parameters can be varied such that impulses in axons of different length arrive at their targets synchronously (Stanford, 1987; Sugihara et al., 1993), suggesting that NOR structure and placement can influence circuit function. Interactions between schwann cell microvilli and the nodal axolemma has been shown to play a critical role in NOR formation and maintenance in the PNS (Eshed et al., 2005). Such microvilli are absent from CNS NOR and it is unclear what nodal component carries out analogous functions in the CNS. Mutations that disrupt nodal structure while preserving myelination lead to decreases in conduction velocity and severe functional deficits (Dupree et al., 1998; Honke et al., 2002; Ishibashi et al., 2002), indicating the importance of defining the exact structural and functional composition of central nodes.

NG2^+^ cells (also termed OPCs or polydendrocytes) are ubiquitous glial progenitors that give rise to oligodendrocytes during development and potentially throughout life (Rivers et al., 2008; Zhu et al., 2008; Kang et al., 2010). NG2^+^ cells form synapses with unmyelinated axons in white matter (Kukley et al., 2007; Ziskin et al., 2007) and have also been shown in electron micrographs to contact NOR in the optic nerve and corpus callosum (Butt et al., 1999; Serwanski et al., 2017), raising the possibility that interaction with axons is a major function of these cells. Immunohistochemical studies have estimated that NG2^+^ cells contact up to 71% of NOR in white matter tracts (Hamilton et al., 2010; Serwanski et al., 2017) and that glycoproteins expressed by spinal cord NG2^+^ cells play a role in limiting neurite outgrowth at NOR (Huang et al., 2005) and can influence conduction of nerve impulses (Hunanyan et al., 2010). If NG2^+^ cells perform critical roles in regulating NOR structure or function, the processes of these glial cells would be expected to constitute a key component of nodal structure. However, detailed quantification of the prevalence of NG2^+^ cell interactions with NOR has not been carried out.

We used transgenic mice in which NG2^+^ cells express membrane-anchored EGFP to examine the spatial relationship between these cells and surrounding axons in 4 CNS white matter tracts. This analysis revealed that the fine processes of these glia are not obligate components of nodal structure. NG2^+^ cells displayed only sparse interaction with surrounding NOR and this interaction was not a key factor guiding NG2^+^ cell process elaboration. In addition, NOR density did not differ between regions with abundant NG2^+^ cells processes and regions devoid of NG2^+^ cell processes. Together, these data suggest that CNS nodes do not require ongoing interaction with NG2^+^ cells and that regulation of nodal structure or function is unlikely to be a dominant functional role of NG2^+^ cells in the adult CNS.

## MATERIALS AND METHODS

### Transgenic mice

*NG2-mEGFP* BAC transgenic mice express membrane-targeted EGFP under control of the NG2 (*Cspg4*) gene promoter. To target EGFP to the plasma membrane, the first 26 amino acids of Lck, a Src family tyrosine kinase which contains two palmitoylation domains and a myristoylation domain, was fused to the N-terminus of EGFP. Both a low *NG2-mEGFP-L* and high *NG2-mEGFP-H* expression line were maintained and used for experiments. Male and female mice were used and all experiments were carried out in strict accordance with protocols approved by the Author’s University.

### Histology

*NG2-mEGFP* mice age P25-P85 were deeply anesthetized with pentobarbital (100 mg/kg) and subjected to cardiac perfusion with 4% paraformaldehyde (PFA) in 0.1 M sodium phosphate buffer in accordance with a protocol approved by the Animal Care and Use Committee at the Author’s Institute. Forebrain (FB), cerebellum (CB), optic nerves (ON), and spinal cord (SC) were gently dissected free and post fixed in 4% PFA at 4°C for 2 hrs. Tissue was cryoprotected in 30% sucrose for 2hrs (ON), 4hrs (SC), or overnight (CB, FB) and then embedded in OTC and sectioned at 50 or 60μm with a cryostat. FB, CB, and SC tissue sections were pre-treated with 2N HCl for 30min at 37°C, followed by 0.1 M sodium borate at room temperature (RT) for 15min. Free-floating sections were then permeabilized/blocked with 0.3% Triton X-100 and 5% normal donkey serum in 0.1 M sodium phosphate buffer for 2 hrs at room temperature. Sections were then incubated with primary antibodies prepared in permeabilizing/blocking solution for 5 hrs at room temperature before being fixed a second time with 4% PFA for 2 hrs at 4°C. Sections were then incubated with secondary antibodies in 5% normal donkey serum in 0.1 M sodium phosphate buffer for 2 hrs at room temperature before mounting on slides. Primary antibodies used: rabbit anti-GFP (1:500, gift from Dr. Huganir, Johns Hopkins), mouse anti-GFP (1:1000, Neuromab), guinea-pig and goat anti-GFP (1:500, gift from Dr. Fukaya, Hokkaido University), mouse pan-sodium channel antibody (1:1000, Sigma), guinea pig anti-Caspr (1:1500, gift from Dr. Bhat, UNC), rabbit and guinea pig anti-NG2 (1:500, gift from Dr. Stallcup, Burnham Institute). Secondary antibodies (raised in donkey): Alexa 488- (Molecular Probes), Cy2-, Cy3-or Cy5-conjugated secondary antibodies to rabbit, mouse, goat or guinea pig (1:500; Jackson ImmunoResearch). Control sections incubated with secondary antibody alone did not result in labeling of cells.

### Image Acquisition and Analysis

Fluorescence images were collected with a LSM 510 Meta confocal microscope (Zeiss). For 3 dimensional reconstruction of NG2^+^ cells and NOR, stacks of confocal images (0.3 μm z-interval) were de-convolved using Autoquant software with a blind deconvolution algorithm. Images were then imported into Imaris image analysis software (Bitplane, RRID:SCR_007370 and SCR_007366 and for reconstruction and quantification. For analysis of individual cells, the filament tracing module was used to reconstruct cell morphology in 3 dimensions and the spots module was used to tag the 3D location of all NOR within the field of view. MatLab XTensions were then used to identify spots (NOR) that fall within different distances of the filament (cell surface). To estimate spatial interactions that occur by chance, the NOR image was flipped about the Y-or Z-axis, or was shifted laterally by 5 μm and analysis was repeated. For analysis of volumes of tissue without focus on individual cells, the surfaces module was used to reconstruct all NG2^+^ surfaces within the field of view. The 3D location of NOR was tagged with the spots module as before and MatLab XTensions were again used to identify NOR within different distances of the reconstructed surface.

### Statistics

Data are expressed as mean ± SEM throughout. Statistical significance when comparing across groups was determined using ANOVA followed by student’s T-test with sequential Bonferroni’s correction for multiple comparisons. A paired t-test was used to determine statistical significance when comparing observed vs. random spatial interactions between NG2^+^ cells and NOR. Details for all statistical tests are listed below.

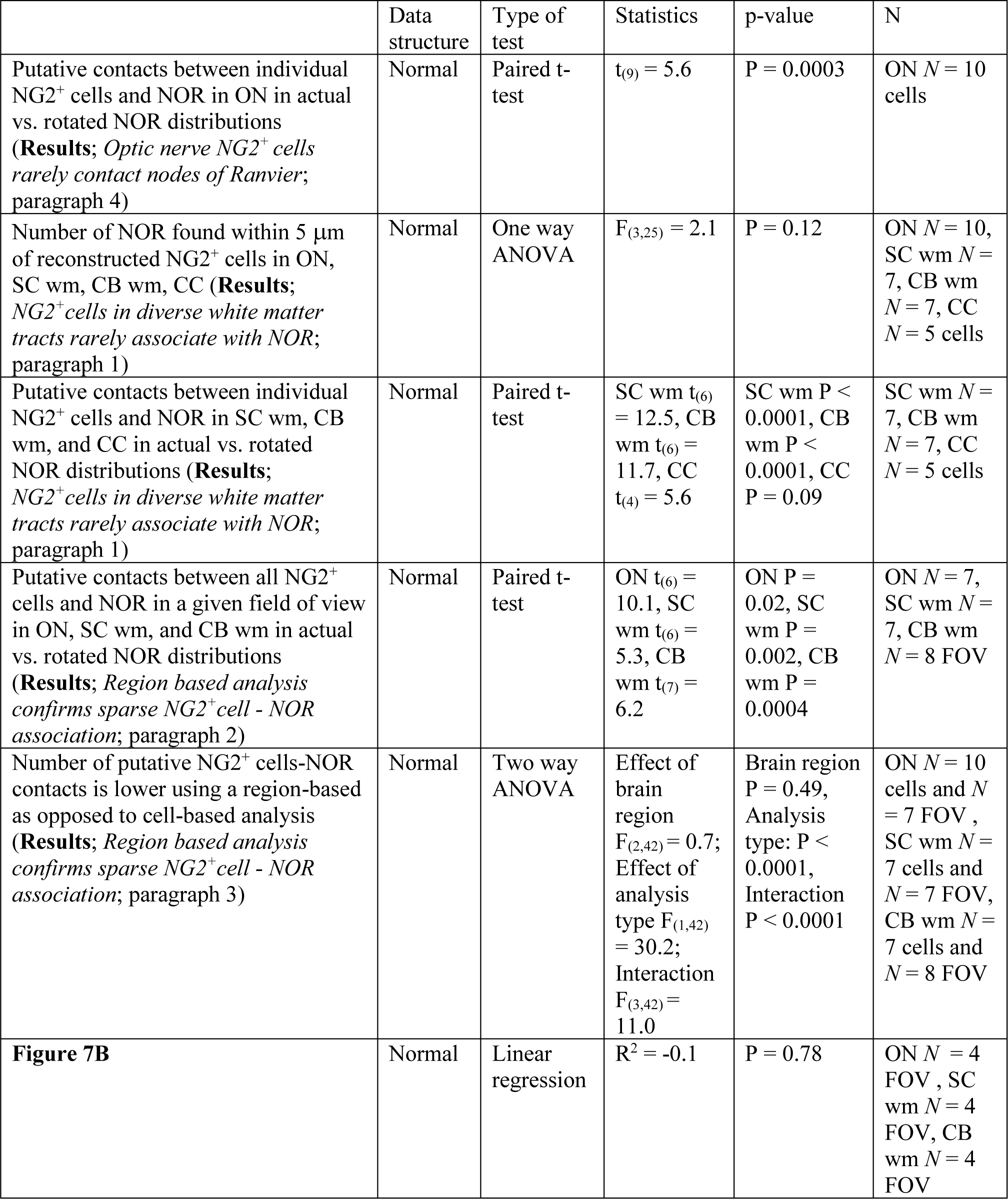
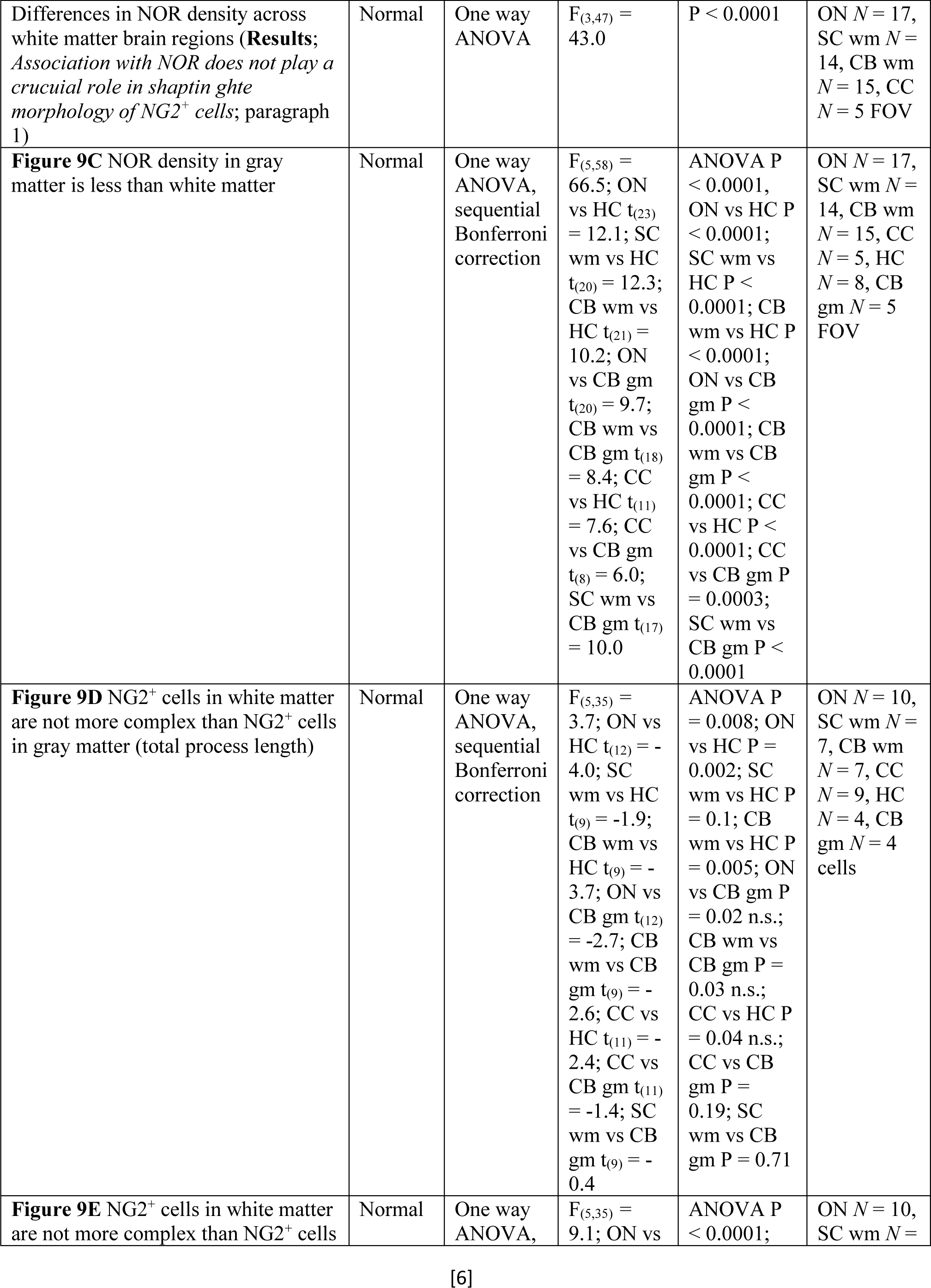
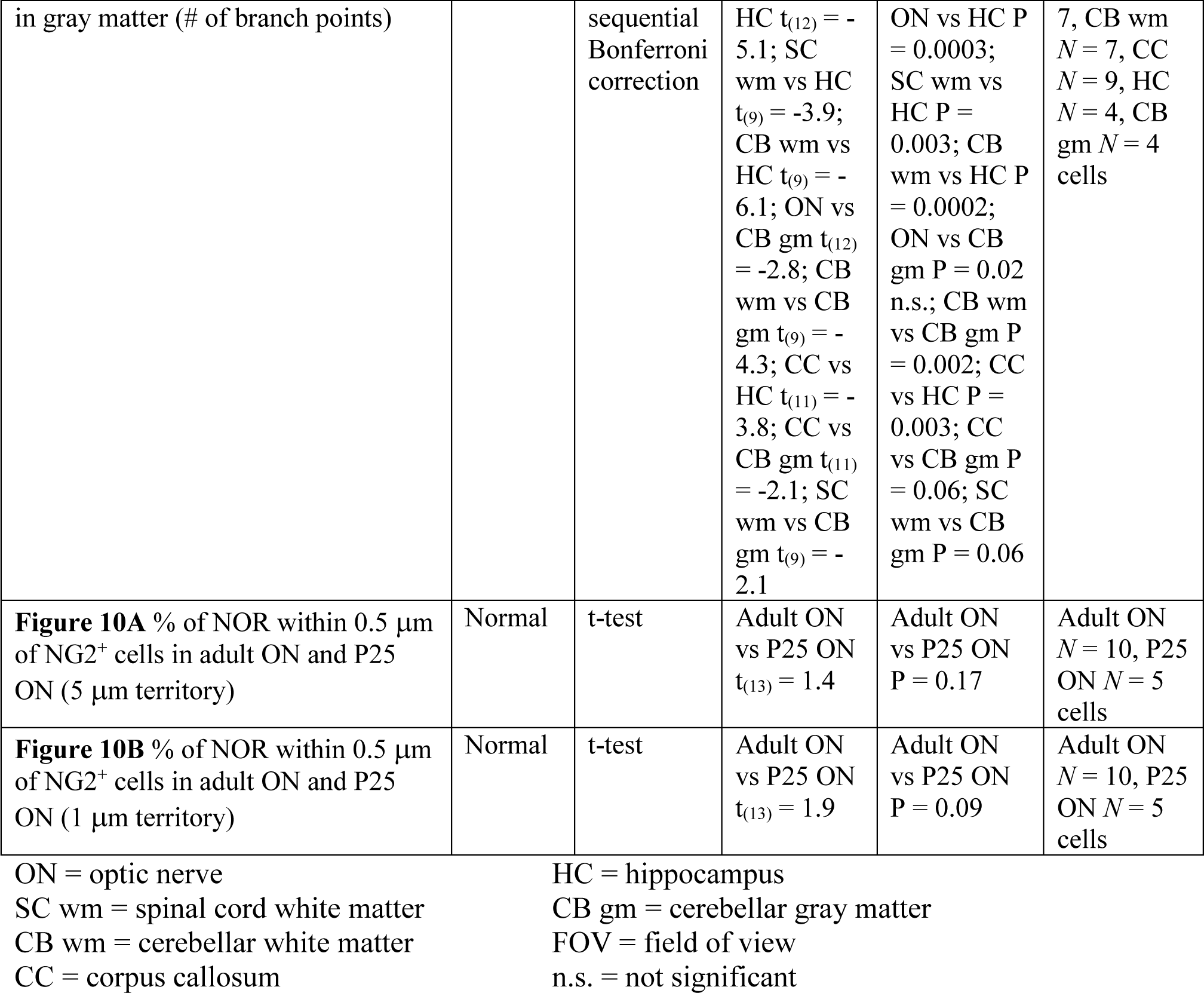

## RESULTS

### Optic nerve NG2^+^ cells rarely contact nodes of Ranvier

To visualize the fine processes of NG2^+^ cells in the CNS, we used BAC transgenic mice that express membrane-targeted EGFP under control of the NG2 promoter (*NG2-mEGFP*) (Figure 1A). In these mice, EGFP^+^ cells are immunoreactive for NG2 (Figure 1A) and PDGF□R, but do not express markers for astrocytes, microglia, or neurons (Hughes et al., 2013). EGFP expression was observed in some pericytes (Figure 1A,B); these endothelial cells, which surround blood vessels, could be unequivocally identified by their bipolar morphology and proximity to blood vessels and were excluded from analysis. Within NG2^+^ cells, EGFP-expression labeled the full extent of distal cell processes and did not alter the number of distal process filopodia (Hughes et al., 2013). Two independent lines of *NG2-mEGFP* mice were maintained, one in which ~90% of NG2^+^ cells express EGFP (high expression line; *NG2-mEGFP-H*) and one in which ~30% of NG2^+^ cells express EGFP (low expression line; *NG2-mEGFP-L*) (Figure 1B,C) (Hughes et al., 2013). In both transgenic lines, EGFP^+^ NG2^+^ cells were distributed evenly throughout gray and white matter regions, suggesting that transgene expression is not limited to a subset of NG2^+^ cells.

**Figure 1.**
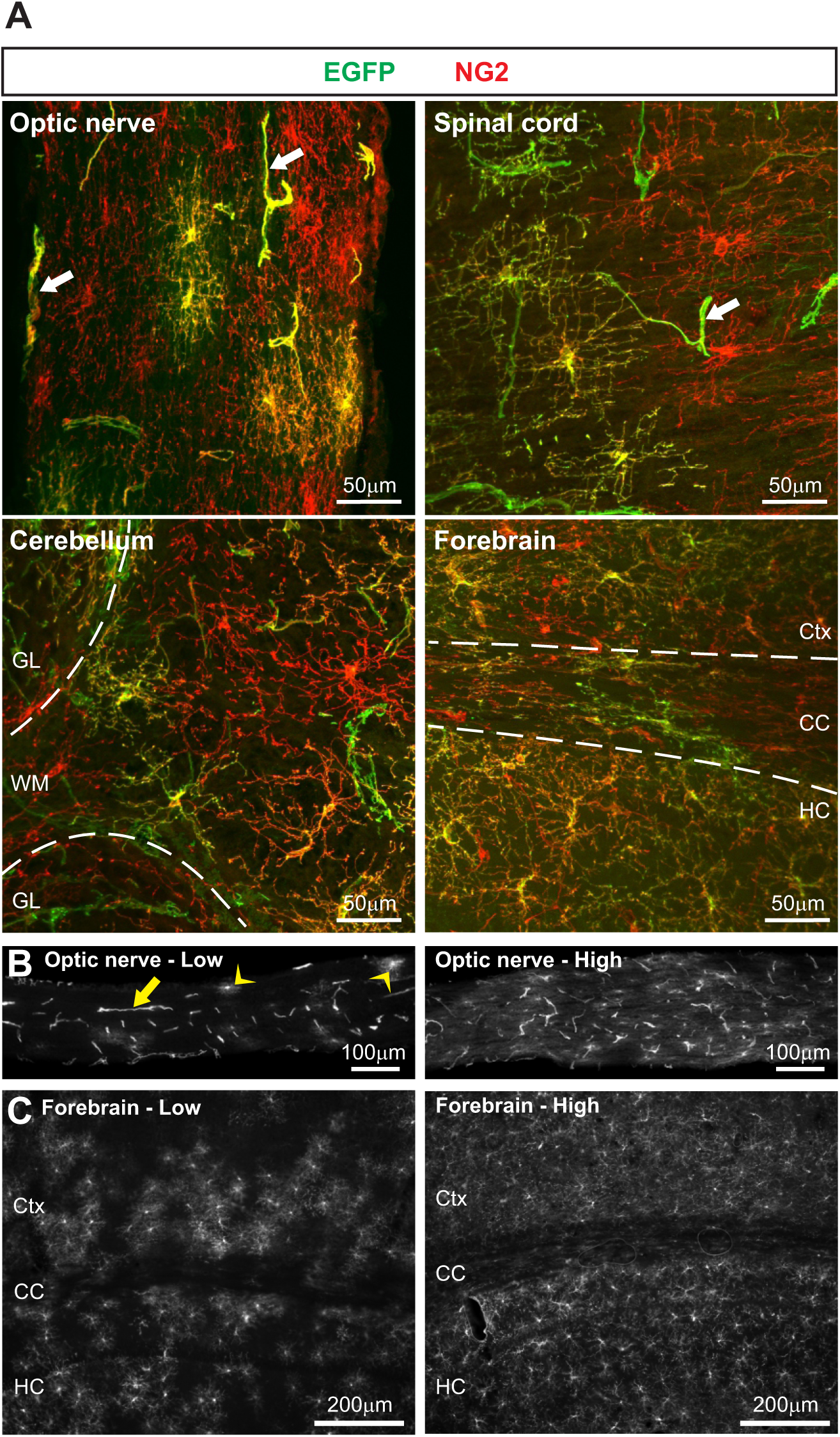
NG2^+^ cells are effectively visualized using *NG2-mEGFP* BAC transgenic mice. ***A***, Immunostaining in representative tissue sections from *NG2-mEGFP* mice demonstrating that EGFP^+^ cells are labeled by antibodies for NG2 in optic nerve, spinal cord white matter, cerebellum, and forebrain. White matter regions in cerebellum and forebrain are outlined with dashed lines. White arrows highlight examples of EGFP^+^ pericytes, which are easily distinguished morphologically and were excluded from analysis. *GL* = granule cell layer, *WM* = white matter, *Ctx* = cortex, *CC* = corpus callosum, *HC* = hippocampus. ***B***, Representative examples of EGFP expression in optic nerve (ON) tissue from low expression (*NG2-mEGFP-L*) and high expression (*NG2-mEGFP-H*) lines of mice. Yellow arrow highlights an EGFP^+^ pericyte and yellow arrowheads highlight two EGFP^+^ NG2^+^ cells. ***C***, Representative examples of EGFP expression in forebrain tissue from *NG2-mEGFP-L* and *NG2-mEGFP-H* mice.

To determine whether individual NG2^+^ cells interact with NOR in their vicinity, we examined EGFP^+^ NG2^+^ cells in optic nerve (ON) from adult (P55-85) *NG2-mEGFP-L* mice. The optic nerve is a uniquely “pure” white matter tract in that it contains axons derived solely from retinal ganglion cells, and 100% of these axons are myelinated. As such, the ON provides an ideal setting for examining the relationship between myelinated axons and the glial cells that surround them. The sparse EGFP expression in *NG2-mEGFP-L* mice allowed unambiguous analysis of isolated EGFP+ NG2^+^ cells (Figure 2A), which displayed thin, branched processes and a roughly bipolar shape oriented along the same axis as nerve fibers, as previously reported (Butt et al., 1999). Stacks of confocal images of individual EGFP^+^ NG2^+^ cells were used to create a 3-dimensional reconstruction of cell morphology (Figure 2A-D). NOR in the surrounding tissue were visualized by immunostaining for voltage-gated sodium channels (NaV), which are clustered at mature nodes, and for Caspr, a cell adhesion protein abundant in paranodal regions flanking NOR (Poliak and Peles, 2003) (Figure 2E,F). NOR were also reconstructed and the 3-dimensional position of each NOR within a confocal image stack was tagged (Figure 2G,H). The accuracy of this positional tagging was confirmed by examining high magnification overlap with NaV and caspr immunostaining (Figure 2I).

**Figure 2.**
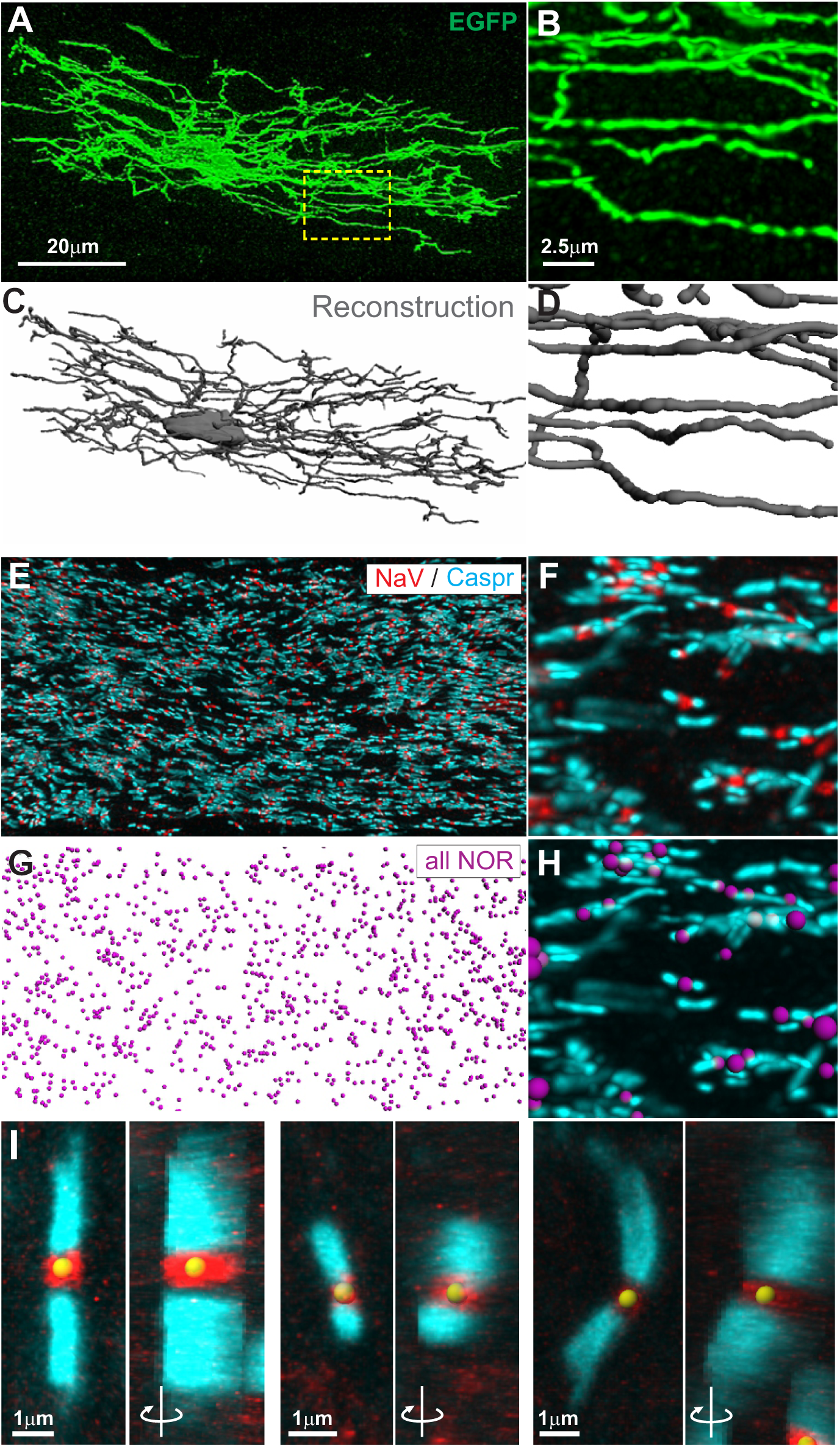
Three dimensional reconstruction of optic nerve NG2^+^ cells and surrounding nodes of Ranvier. ***A***, Maximum intensity projection of an isolated, EGFP-expressing NG2^+^ cell in optic nerve tissue from an *NG2-mEGFP-L* mouse (P56). *Dashed box* shown at higher magnification in ***B. C and D***, 3D reconstruction of cell morphology. ***E***, Nodes of Ranvier (NOR) in the same field of view as *A* revealed by immunostaining for voltage-gated sodium channels (NaV) in red and paranodal protein caspr in cyan; enlarged view shown in ***F. G and H***, 3D tagging of NOR shown together with caspr immunostaining in cyan. ***I***, Three examples of NOR with NaV (red) and caspr (cyan) immunostaining illustrating the accuracy of NOR tagging (yellow sphere); panels on the right are rotated 90° to show the x-z dimension.

As highlighted both by immunostaining and 3D positional tagging, NOR are found at very high density within optic nerve tissue. To determine whether NG2^+^ cells interact with NOR in their vicinity, we first defined which NOR fall within the “territory” of an individual reconstructed cell. NG2^+^ cells exhibit a tiled distribution and are not normally observed to extend processes into regions occupied by neighboring NG2^+^ cells (Dawson et al., 2003; Wigley and Butt, 2009). For this reason, NOR within 5 μm of the reconstructed cell were taken as a reasonable estimation of the territory of that cell (Figure 3A). For 10 reconstructed optic nerve NG2^+^ cells, there were 619 ± 45 NOR within 5 μm of NG2^+^ cell processes, consistent with the high density of NOR in this white matter tract. However, only a small percentage of these NOR (14 ± 2 %, *N* = 10) were found within 0.5 μm of any portion of the NG2^+^ cell membrane (Figure 3A-C). This percentage is likely an overestimation of the instances of actual contact between NG2^+^ cells and NOR, as electron microscopic analysis indicates that NG2^+^ cell processes interacting with NOR come within nanometers of the exposed nodal axolemma (Butt et al., 1999).

**Figure 3.**
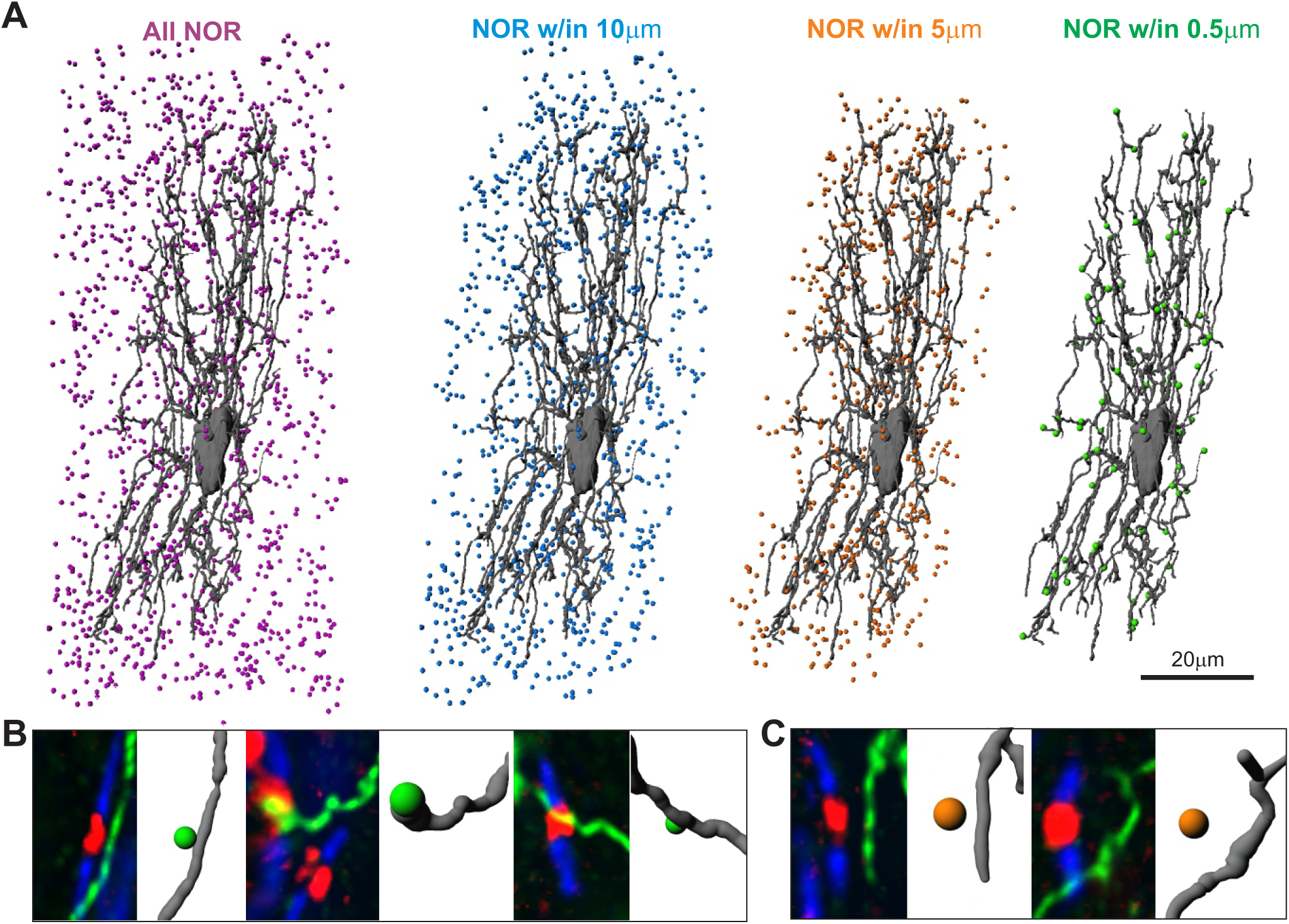
NG2^+^ cell territory and close associations with NOR. ***A***, The reconstructed NG2^+^ cell shown in Fig. 2 and tagged NOR from the entire field of view (purple), or only those NOR that come within 10 μm (blue), 5 μm (orange), or 0.5 μm (green) of the cell surface. ***B***, Three examples of NOR that came within 0.5 μ of the cell surface; immunostaining for Nav (red), caspr (blue), and EGFP (green) shown at *left* and 3D reconstruction of the cell surface and tagged NOR shown at *right*. ***C***, Two examples of NG2^+^ cell terminal processes that pass by but do not come within 0.5 μm of surrounding NOR.

Because the density of axons is high within optic nerve, many NOR are likely to be in close proximity to the highly ramified processes of NG2^+^ cells simply by chance. To estimate the potential number of random spatial interactions, the NOR distribution for a given cell was rotated 180 degrees about the y-axis and quantification was repeated. With this approach 7 ± 1% of NOR within the territory of an NG2^+^ cell came within 0.5 μm of the cell surface, indicating that approximately half of the observed associations can be explained by chance. Similar analysis was carried out by shifting NOR distributions laterally 5 μm or rotating NOR distributions about the Z-axis and comparable estimates of random associations were obtained (Table 1). This analysis indicates that > 90% of NOR within the territory of individual optic nerve NG2^+^ cells are not associated with NG2^+^ cell processes. However, the number of putative interactions observed with the actual NOR distribution was significantly greater than the number of associations expected by chance (P = 0.0003, paired t-test), suggesting that NG2^+^ cells may form *bona fide* contacts with a small portion of NOR in their vicinity.

**Table 1.**
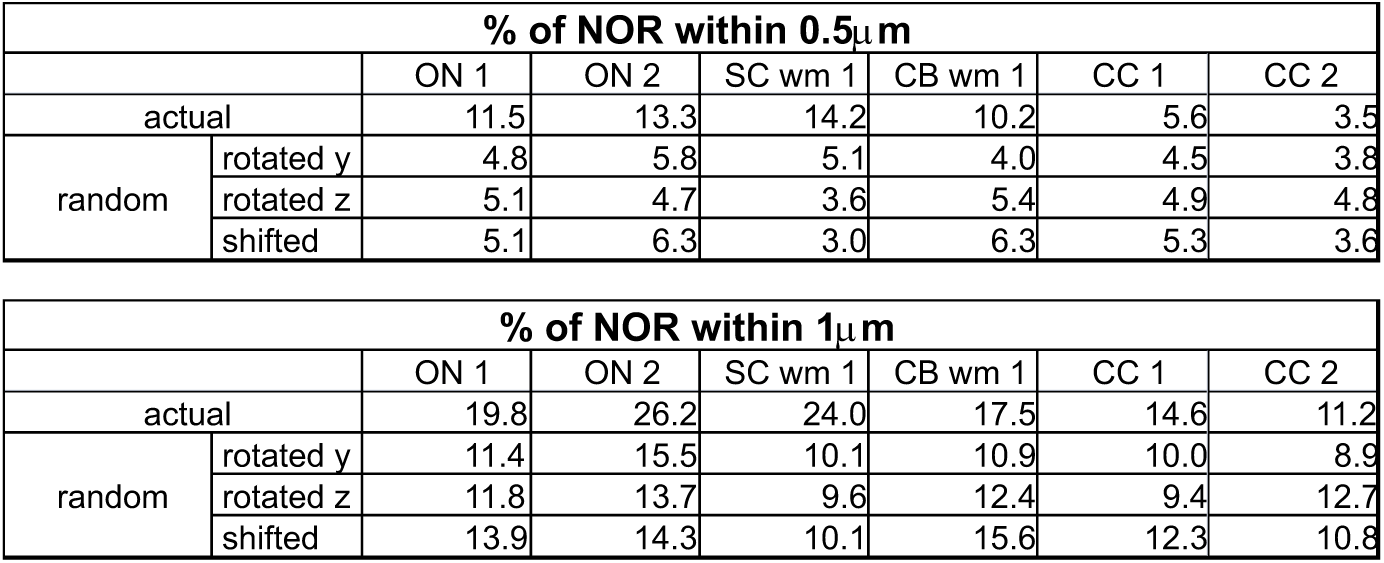
Estimation of random spatial interactions. For six example NG2 ^+^ cells, the % of NOR that came within 0.5 or 1 μm of the cell surface in the actual NOR distribution is listed. Random spatial interactions were estimated by rotating NOR distributions 180 degrees about the Y-or Z-axis or shifting the distribution laterally by 5 μm and re-calculating the % of NOR found within 0.5 of 1 μm of the cell surface.

### NG2^+^ cells in diverse white matter tracts rarely associate with NOR

In spinal cord white matter, NG2^+^ cells have been implicated in limiting neurite outgrowth at NOR either by direct contact with NOR or through deposition of Oligodendrocyte Myelin Glycoprotein (OMgp) (Huang et al., 2005). In addition, interaction of the NG2 proteoglycan with spinal cord NOR has been suggested to alter nerve impulse conduction (Hunanyan et al., 2010). Together, these findings suggest that NG2^+^ cells may play a key role in regulating the function of spinal cord NOR, a conclusion that would seem at odds with the very sparse NG2^+^ cell - NOR interaction observed in the ON. To determine whether the limited interaction between NG2^+^ cells and NOR observed in ON is a consistent feature of CNS white matter tracts, we performed similar analysis in spinal cord white matter (SC wm), cerebellar white matter (CB wm), and the corpus callosum (CC) (Figure 4). These white matter tracts are more heterogeneous than ON, containing both myelinated and unmyelinated axons that arise from diverse neuronal populations. However, nodal structure is relatively consistent across white matter regions and NG2^+^ cells are ubiquitous throughout the CNS. Despite the increased heterogeneity of these white matter regions, the number of NOR found within 5 μm of reconstructed NG2^+^ cells did not differ significantly from that observed in ON (ON 619 ± 45 *N* = 10, SC wm 703 ± 54 *N* = 7, CB wm 537 ± 50 *N* = 7, CC 563 ± 32 *N* = 5; P = 0.12, ANOVA). Consistent with the analysis in ON, only a small percentage of NOR in the territory of NG2^+^ cells in these additional white matter regions came within 0.5 μm of the cell surface (SC wm 14 ± 1%, CB wm 11 ± 1%, CC 7 ± 2%) (Figure 4B). Random instances of close NG2^+^ cell-NOR proximity were estimated as before (SC wm 5 ± 1%, CB wm 4 ± 0.3%, CC 4 ± 1%) and in all regions except CC, the number of NOR within 0.5 μm was significantly greater than what would be expected due to chance (SC wm P = 0.000001, CB wm P = 0.00002, CC P = 0.08; paired t-test). These values were then used to derive a “corrected” estimate of functionally relevant spatial interactions between NG2^+^ cells and surrounding NOR (ON 7.0 ± 1%, SC wm 8.9 ± 0.5%, CB wm 7.1 ± 1%, CC 3.0 ± 1%)(Figure 4B). Together, these data indicate that the spatial relationship of individual NG2^+^ cells with NOR in the surrounding tissue is consistent across diverse CNS white matter tracts and that NG2^+^ cells are in contact with only a small percentage of NOR.

**Figure 4.**
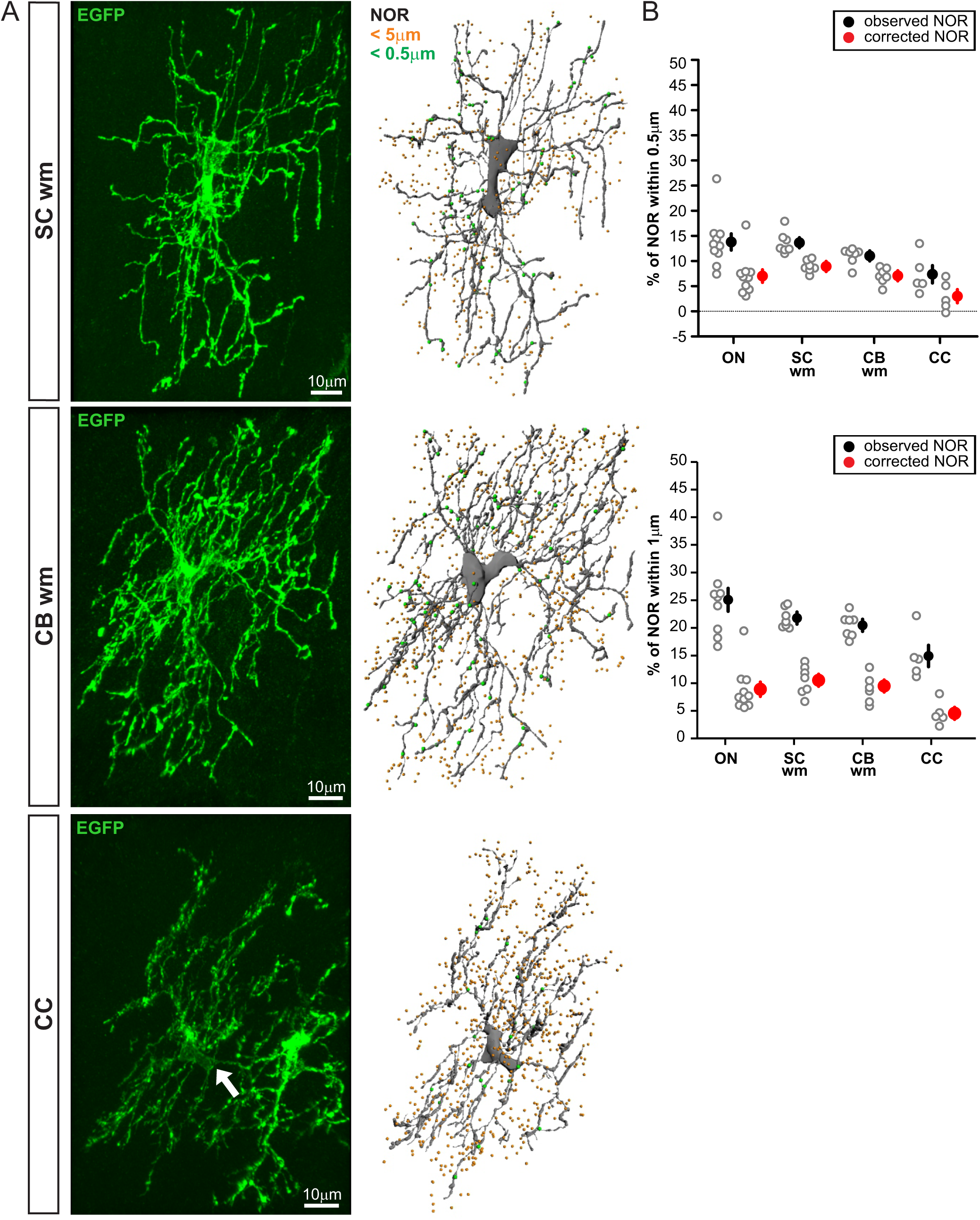
Sparse NG2^+^ cell-NOR interaction is consistent across diverse white matter tracts. ***A***, Immunostaining for representative EGFP^+^ NG2^+^ cells in spinal cord white matter (SC wm), cerebellar white matter (CB wm), and corpus callosum (CC) *at left*. (Two EGFP+ NG2^+^ cells are visible in the corpus callosum; reconstructed cell indicated by *white arrow*). 3D reconstruction for each NG2^+^ cell shown *at right*. NOR within 5 μm of the cell are shown in orange and those in close proximity to the cell surface (within 0.5 μ) are shown in green. ***B***, Percentage of NOR found within 0.5 μm of the cell surface from the pool of NOR that come within 5 μm of the cell (*top*). Percentage of NOR found within 1 μm of the cell surface from the same pool of NOR that are within a 5 μm territory of the cell. *Red circles* show the number of spatial interactions after correcting for interactions expected due to chance (% of NOR within 0.5 μm using actual NOR distribution - % of NOR within 0.5 μm using NOR distribution flipped about the Y-axis).

### Region based analysis confirms sparse NG2^+^cell - NOR association

Morphological reconstruction of individual NG2^+^ cells facilitates analysis of where along NG2^+^ cell processes putative interactions occur and whether subsets of NG2^+^ cells interact with NOR to differing degrees. However, this approach is limited by the necessity of defining the spatial territory of reconstructed cells. To determine whether the particular parameters of this analytical approach affect the detection of NG2^+^ cell – NOR interactions, the definition of “proximity” was relaxed to 1 μm. As expected, in this analysis a higher percentage of NOR came within 1 μm of the cell surface (ON 25 ± 2% *N* = 10, SC wm 22 ± 1% *N* = 7, CB wm 20 ± 1% *N* = 7, CC 15 ± 2% *N* = 5) (Figure 4B). However, the number of interactions estimated to occur by chance rose as well (ON 16 ± 1%, SC wm 11 ± 1%, CB wm 11 ± 1%, CC 10 ± 1%), leading to the same conclusion that putative interactions occur between NG2^+^ cells and a small percentage of NOR in their vicinity (ON 9 ± 2%, SC wm 11 ± 1%, CB wm 10 ± 1%, CC 5 ± 1%; observed%-random%). Similarly, when analysis was repeated using a cell “territory” of 10 μm or 2 μm, results did not differ significantly (Figure 5A,B).

**Figure 5.**
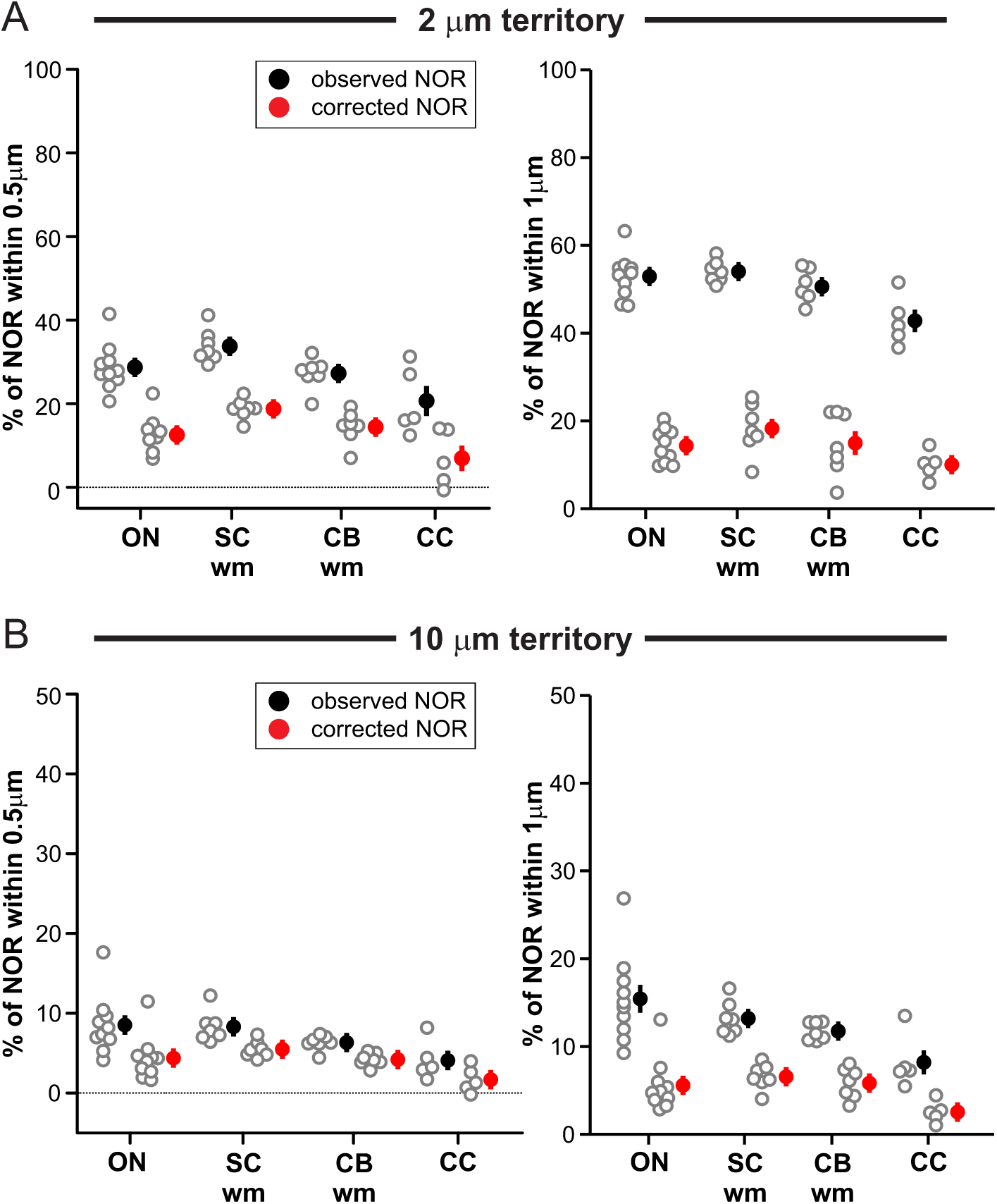
Analysis of NG2^+^ cell proximity to NOR using different definitions of cell territory. ***A***, Percentage of NOR that come within 0.5 μm (*left*) or 1.0 μm (*right*) of NG2^+^ cell processes calculated from the pool of NOR that fall within 2.0 μm of the cell (*black circles*) and the number of interactions after correcting for interactions expected due to chance (*red circles*). ***B***, Percentage of NOR that come within 0.5 μm (*left*) or 1.0 μm (*right*) of NG2^+^ cell processes calculated from the pool of NOR that fall within 10.0 μm of the cell. ON = optic nerve, SC wm = spinal cord white matter, CB wm = cerebellar white matter, CC = corpus callosum.

As a final confirmation that analysis of individual NG2^+^ cells accurately reflects the relationship of these glia with surrounding NOR, we used the high expression *NG2-mEGFP-H* line of mice to quantify the proximity of NOR and NG2^+^ cell processes in larger volumes of white matter without focusing on individual cells (Figure 6). In a given region of interest, all NG2^+^ cell processes were reconstructed in 3 dimensions and the 3D position of NOR was tagged as before. Of all NOR present in the examined regions, only a small percentage came within 0.5 μm of an NG2^+^ cell surface (ON 13 ± 3% *N* = 7, SC wm 4 ± 0.5% *N* = 7, CB wm 4 ± 0.2% *N* = 8) (Figure 6B). Consistent with the analysis of individual NG2^+^ cells, the observed number of associations was significantly greater than would be expected due to chance alone (ON P = 0.02, SC wm P = 0.002, CB wm P = 0.0004, paired t-test; random associations ON 9 ± 2%, SC wm 2 ± 0.2%, CB wm 2 ± 0.3%, rotated NOR distribution). Corpus callosum was not included in this analysis because EGFP was not expressed by all callosal NG2^+^ cells in this line of mice. When the definition of physical proximity was relaxed to 1 μm (Figure 6C), the number of NOR associated with NG2^+^ cell processes increased, but so did the number of estimated random interactions, leading to a similar conclusion that only a small percentage of NOR within a given volume of tissue may possess *bona fide* associations with NG2^+^ cell processes (ON 5 ± 2%, SC wm 2 ± 0.4%, CB wm 2 ± 0.3%; actual%-random%). Together these data indicate that analysis of individually reconstructed NG2^+^ cells is a valid approach for probing their relationship with surrounding NOR.

**Figure 6.**
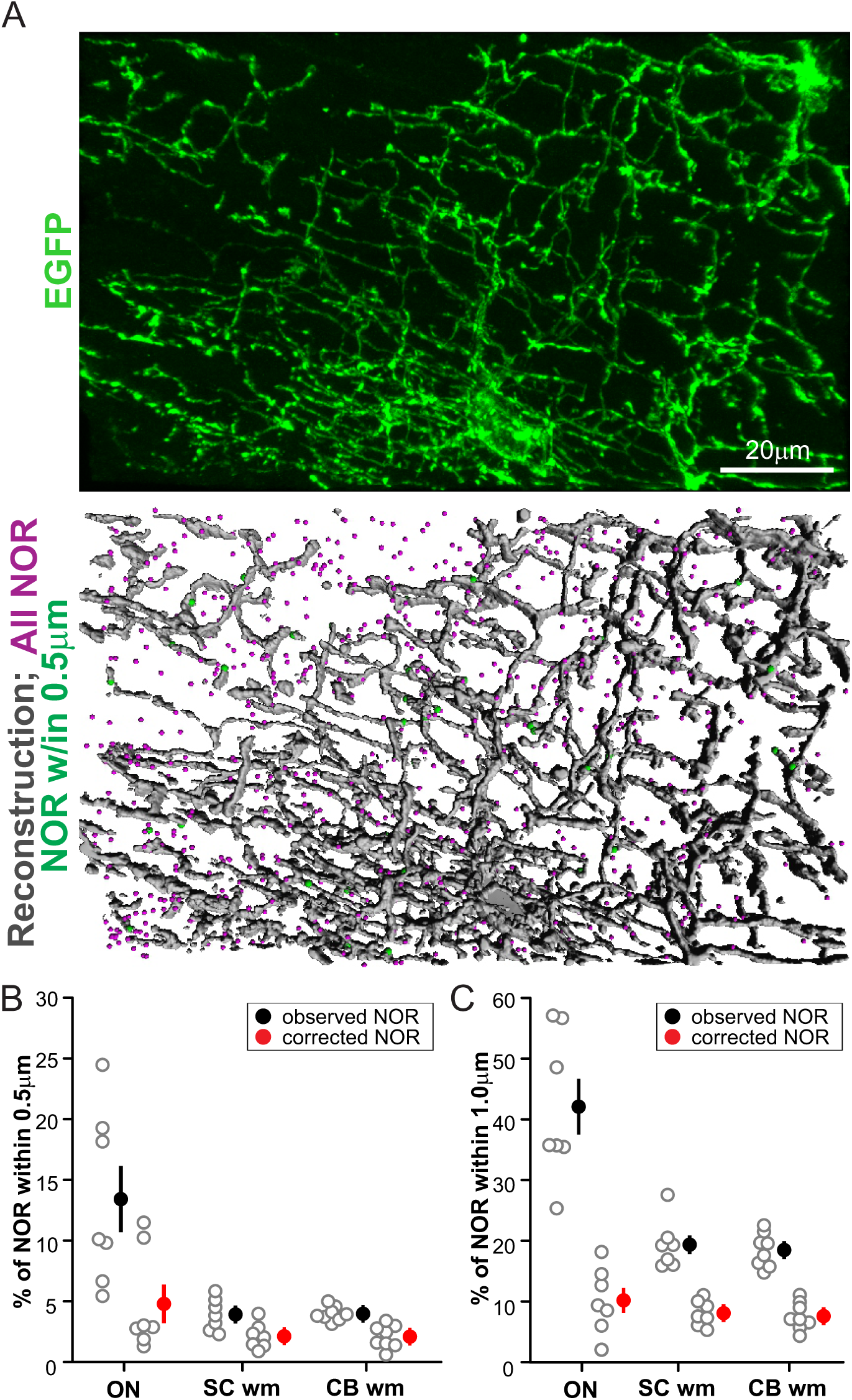
Region based analysis confirms sparse NG2^+^ cell-NOR interaction. ***A***, EGFP expression (*top*) and 3D reconstruction of all EGFP^+^ processes (*bottom*) in a volume of spinal cord white matter (SC wm). All NOR found within the region are colored *magenta*; NOR that come within 0.5 μm of NG2^+^ cell processes are colored in *green*. ***B***, Percentage of all NOR within analyzed tissue volume that come within 0.5 μm of NG2^+^ cell processes. *Red circles* show the number of interactions after correcting for interactions expected due to chance. ***C***, Percentage of all NOR within analyzed tissue volume that come within 1.0 μm of NG2^+^ cell processes. *Red circles* show the number of interactions after correcting for interactions expected due to chance. ON = optic nerve, CB wm = cerebellar white matter.

One key difference that emerged between region-based analyses and data from individually reconstructed NG2^+^ cells, is that the number of observed putative interactions between NG2^+^ cells and NOR was significantly lower using a region-based approach (P < 0.0001, Two-way ANOVA). Immunostaining for NG2 or examination of *NG2-mEGFP-H mice* reveals small zones within white matter that are devoid of NG2^+^ cell processes (Figure 6A and 7A). These “voids” are not taken into consideration when quantifying only NOR that fall within the territory of an individual cell; hence, this cell-based quantification approach could lead to an overestimation of the prevalence of NG2^+^ cell-NOR associations. NOR density was comparable in regions with sparse or abundant NG2^+^ cell processes, and no correlation was observed between the density of NOR and the degree of NG2^+^ cell process coverage (Figure 7B), supporting the conclusion that NG2^+^ cell processes are not requisite members of NOR whose continual presence is required for ongoing NOR function or maintenance in the adult CNS.

**Figure 7.**
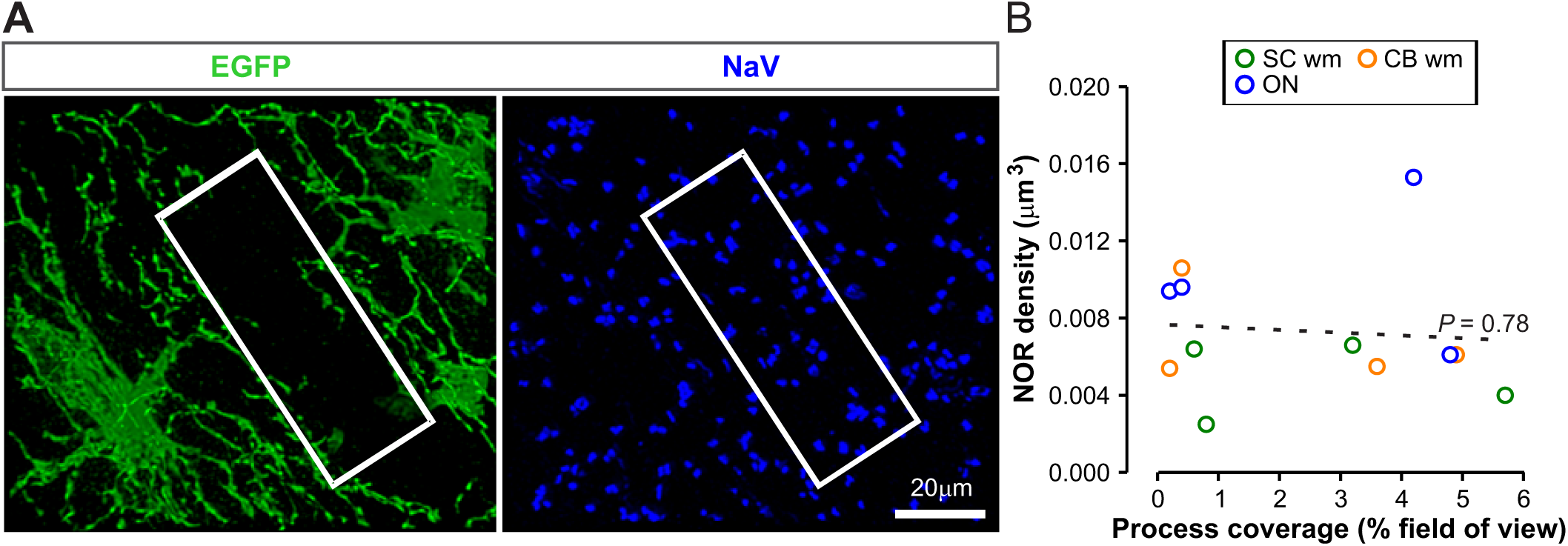
NOR density is unchanged in regions devoid of NG2+ cell processes. ***A***, Immunostaining in spinal cord white matter from P56 high expression *NG2-mEGFP-H* mouse highlighting a region in which NG2^+^ cell processes are largely absent (*white box*). ***B***, NOR density was quantified in optic nerve (ON), spinal cord white matter (SC wm), and cerebellar white matter (CB wm) and plotted against the degree to which the field of view was covered by NG2^+^ cell processes; no correlation was observed between the density of NOR and the degree of NG2^+^ cell process coverage (R^2^ = −0.1, P = 0.78).

### NG2^+^ cell processes rarely terminate at NOR

Electron microscopic approaches indicate that NG2^+^ cell processes which contact NOR appear to enwrap the exposed axolemma (Butt et al., 1999), suggesting that NG2^+^ cells may extend specialized processes which terminate at NOR. Our data support the idea that NG2^+^ cells interact with a small percentage of NOR in the surrounding neuropil. However, among the NOR that are within 0.5 μm of NG2^+^ cell surfaces, the majority were found along cell processes (*en passant*) (ON 76 ± 2% *N* = 10, SC wm 74 ± 2% *N* = 7, CB wm 77 ± 2% *N* = 7, CC 88 ± 2% *N* = 5) rather than at the tips of NG2^+^ cell processes (ON 24 ± 2%, SC wm 26 ± 2%, CB wm 23 ± 2%, CC 14 ± 2%)(Figure 8A,B). In addition, NOR associations with NG2^+^ cell process tips can also be observed in the random NOR distribution (% of NOR w/in 0.5 μm found at process tips; ON 18 ± 2%, SC wm 12 ± 2%, CB wm 13 ± 1%, CC 12 ± 2%)(Figure 8A), indicating that some of these apparent tip-NOR interactions may be due to chance alone. Finally, only a small subset of NG2^+^ cell process tips terminated at NOR (% of tips w/in 0.5 μm of NOR; ON 8 ± 1% *N* = 10, SC wm 8 ± 1% *N* = 7, CB wm 9 ± 1% *N* = 7, CC 3 ± 1%) (Figure 8C). Together these data suggest that attraction to NOR is not the primary factor guiding the extension of NG2^+^ cell process tips.

**Figure 8.**
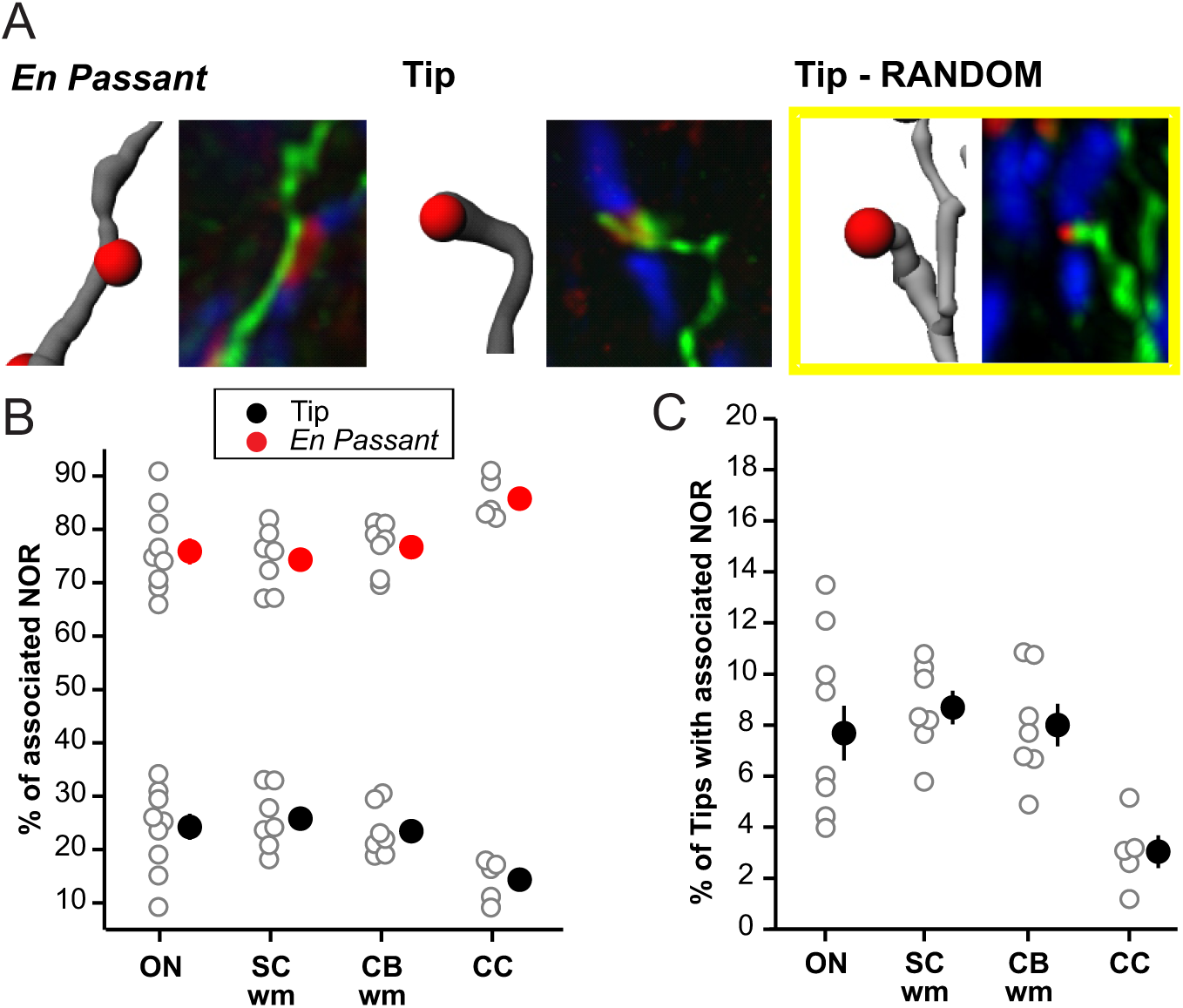
NG2^+^ cell processes rarely terminate at NOR. ***A***, Examples of close proximity NOR (located within 0.5 μm of cell) that were found *en passant* along NG2^+^ cell processes, or at the tips of NG2^+^ cell processes; *yellow box* highlights NG2^+^ cell-NOR interaction that occurred by chance (*random*). ***B***, Percentage of close proximity NOR (located within 0.5 μm of cell) that were found *en passant* along NG2^+^ cell processes (*red circles*) vs. at the tips of NG2^+^ cell processes (*black circles*). ***C***, Percentage of NG2^+^ cell process tips that terminate near (within 0.5 μ) NOR.

### Association with NOR does not play a crucial role in shaping the morphology of NG2^+^ cells

To further explore whether interactions with NOR play a key role in regulating NG2^+^ cell process extension, we performed detailed morphological comparisons of gray and white matter NG2^+^ cells. If NOR interaction played a dominant role in regulating NG2^+^ cell process extension, then NG2^+^ cells in regions of high NOR density (i.e. white matter tracts) should be morphologically more complex than NG2^+^ cells in regions of low NOR density (i.e. gray matter regions). To test this hypothesis, individual NG2^+^ cells in two gray matter regions, CA1 hippocampus (HC) and cerebellar molecular layer (CB ml), were morphologically reconstructed as described above (Figure 9A) and compared to individually reconstructed white matter NG2^+^ cells. Gray matter regions displayed dramatically lower NOR density compared to white matter regions (P < 0.0004 all gray matter vs. white matter region comparisons, t-test) (Figure 9C). While CA1 HC contained a small number of identifiable NOR, CB ml showed only diffuse labeling with NaV antibody and a complete absence of NOR (Figure 9B). This observation is consistent with existing literature indicating that parallel fibers in the rodent CNS are unmyelinated (Wyatt et al., 2005) and NOR and axon initial segments were clearly visible in the adjacent Purkinje and granule cell layers, indicating that immunolabeling for NOR is effective in cerebellar tissues. Despite a dramatically lower NOR density, gray matter NG2^+^ cells displayed highly complex morphologies (Figure 9D,E). Total process length of gray matter NG2^+^ cells was comparable to or exceeded that of white matter NG2^+^ cells (ON vs HC P = 0.002; CB wm vs HC P = 0.005; all other white matter vs. gray matter comparisons n.s., t-test). Similarly, number of branch points of gray matter NG2^+^ cells was comparable to or exceeded that of white matter NG2^+^ cells (ON vs HC P = 0.0003; SC wm vs HC P = 0.003; CB wm vs HC P = 0.0002; CB wm vs CB gm P = 0.002; CC vs HC P = 0.003; all other white matter vs. gray matter comparisons n.s., t-test). Together, these observations support the conclusion that interaction with NOR does not play a dominant role in determining NG2^+^ cell arborization.

**Figure 9.**
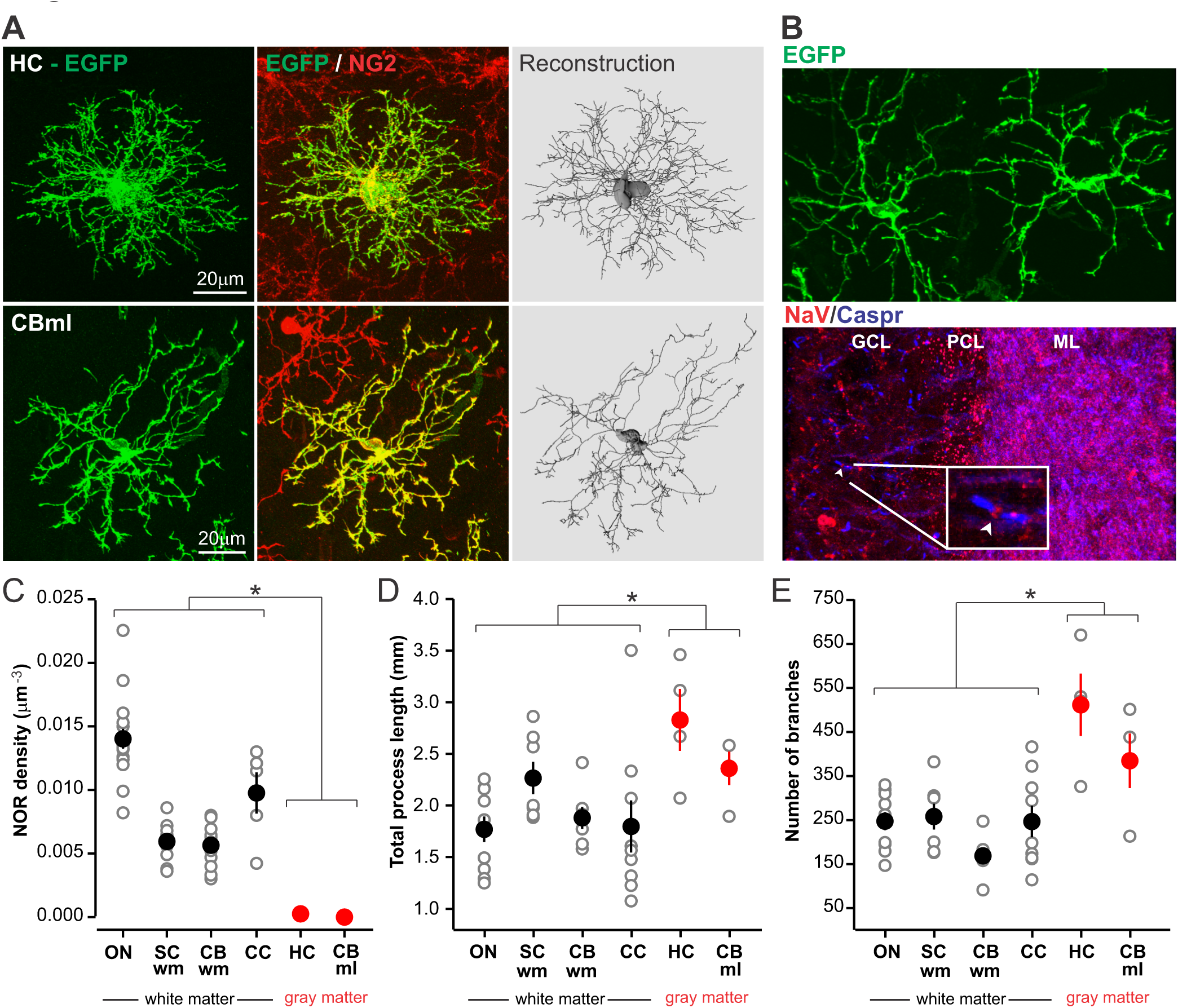
Attraction to NOR does not significantly influence NG2^+^ cell morphology. ***A***, Immunostaining and 3D reconstruction of individual NG2^+^ cells in CA1 region of hippocampus (HC) and the molecular layer of cerebellum (CB ml). ***B***, EGFP+ NG2^+^ cells in cerebellum with very similar morphology (*top panel*) despite the absence of NOR in the molecular layer (ML) and presence of NOR in the granule cell layer (GCL) (*bottom panel*); *arrowhead* and *inset* highlight a well-defined NOR within the granule cell layer; PC = purkinje cell layer. ***C***, Density of NOR in all analyzed brain regions highlighting the difference in density between white matter regions (*black circles*) and gray matter regions (*red circles*) * P < 0.0004 all gray matter vs. white matter region comparisons, t-test. ***D***, Total process length for white matter (*black circles*) and gray matter (*red circles*) NG2^+^ cells. * P < 0.006 ON vs HC, CB wm vs HC; all other white matter vs. gray matter comparisons n.s., t-test ***E***, Total number of branch points for white matter (*black circles*) and gray matter (*red circles*) NG2^+^ cells. * P < 0.004 ON vs HC, SC wm vs HC, CB wm vs HC, CB wm vs CB gm, CC vs HC; all other white matter vs. gray matter comparisons n.s., t-test.

## DISCUSSION

Nervous system function depends on the successful propagation of nerve impulses over long distances. The critical role that myelination and NOR play in such long range signaling has been appreciated since electrophysiology studies in the late 1940’s (Huxley and Stampfli, 1949). The NOR is a highly specialized structure with elaborate adhesive and signaling interactions between the axon and surrounding glial cells. In the PNS, Schwann cell filopodia play critical roles in node formation and maintenance. In the CNS, interactions between myelinating oligodendrocytes and the underlying axon support the proper segregation of voltage-gated sodium and potassium channels to nodal and perinodal regions, respectively (Poliak and Peles, 2003). The processes of perinodal astrocytes contact the exposed nodal axolemma and are thought to play key roles in maintaining local ion homeostasis (Waxman et al., 1994). However, numerous questions about the formation and maintenance of CNS NOR persist, and it is unclear which nodal elements play functional roles analogous to the Schwann cell filopodia in the PNS.

NG2^+^ cells arise from bipolar progenitors which migrate outward from the sub-ventriclular zone during late embryogenesis (Richardson et al., 2006). These NG2^+^ progenitors then elaborate numerous branched processes and ultimately populate the entire CNS. During early postnatal periods, NG2^+^ cells give rise to oligodendrocytes but remain ubiquitous throughout life (Levison et al., 1999; Rivers et al., 2008). While NG2^+^ cells could serve as an oligodendrocyte reservoir to support ongoing myelin turnover or remyelination following a demyelinating insult, the role that these progenitors play in the adult CNS is still unclear. BrdU labeling studies suggest that a subset of adult NG2^+^ cells is quiescent (Psachoulia et al., 2009), raising the possibility that these non-proliferative NG2^+^ cells have unique functional roles. Contact between NG2^+^ cells and NOR has been clearly observed or inferred using electron microscopic approaches in optic nerve of adult rats (Butt et al., 1999; Serwanski et al., 2017) and spinal cord of mice (Serwanski et al., 2017), suggesting that some NG2^+^ cells may participate in regulating nodal function.

Immunostaining for NG2 proteoglycan yielded high estimates of the percent of NOR contacted by NG2 cell processes (33-71%) (Hamilton et al., 2010; Serwanski et al., 2017) and NG2^+^ cell processes in the spinal cord white matter have been suggested to regulate the formation of axon collaterals at NOR (Huang et al., 2005) and to influence conduction efficacy (Hunanyan et al., 2010). These findings invite a revised understanding of nodal structure and function in which NG2^+^ glia play a key role in shaping neuronal impulses, analogous to the tripartite synapse model in which glia actively modulate signaling at central synapses (Perea et al., 2009). However, the prevalence of NG2^+^ cell processes at NOR has not been rigorously quantified in the adult CNS, nor have the instances of apparent contact in light microscopic analysis that occur by chance alone been estimated.

In the present study, we used transgenic and light microscopic approaches to obtain the first comprehensive quantification of the spatial relationship between NG2^+^ cells and NOR in the adult CNS. To effectively visualize the highly ramified processes of NG2^+^ cells, we used BAC transgenic mice that express membrane-targeted EGFP under control of the NG2 promoter (*NG2-mEGFP*). In these mice, EGFP expression is restricted to NG2^+^ cells, is distributed throughout the entire cell membrane, as assessed by immunostaining for GFP and NG2, and provides superior labeling of cell processes (Hughes et al., 2013). Because NG2^+^ cells are tiled throughout the CNS, a particular advantage of the *NG2-mEGFP-L* low expression line was the ability to clearly distinguish the processes of an individual NG2^+^ cell from those of neighboring NG2^+^ cells, which is challenging with immunostaining. While light microscopy cannot approach the nanometer resolution offered by electron microscopic approaches, acquisition of stacks of confocal images from *NG2-mEGFP-L* mice allowed us to perform quantitative analysis of entire cells and determine where along NG2^+^ cell process putative interactions with NOR occur. Our data are consistent with the idea that interaction between NG2^+^ cells and NOR does take place. However, using several approaches, we find that in 4 distinct CNS white matter tracts a minority of NOR (7-14%) are in close proximity to NG2^+^ cell processes. Ultrastructural analysis indicates that astrocytes and NG2^+^ cells that contact NOR come within nanometers of the exposed axolemma, suggesting that our analysis likely represents an overestimation of the instances of actual contact between these structures. These data indicate that NG2^+^ cells are not a permanent fixture at the majority of NOR, making it unlikely that they regulate neurite outgrowth or influence conduction efficacy at these structures. The observation of pockets of white matter devoid of NG2^+^ cell processes where NOR density is unaltered also argues that the continuous presence of NG2^+^ cell processes is not necessary for proper node functioning.

Despite the fact that NG2^+^ cell processes are not present at most NOR, the number of observed interactions between NG2^+^ cells and NOR was greater than would be expected due to chance alone, suggesting that NG2^+^ cells do associate with a small number of NOR in their vicinity. Though regional differences in NG2^+^ cell morphology have been noted qualitatively (Dawson et al., 2003), the factors governing elaboration of NG2^+^ cell processes have not been demonstrated. In addition, quantitative analysis of NG2^+^ cell morphology has been very limited. Does attraction to NOR influence the elaboration of NG2^+^ cell processes? Though the density of NOR is dramatically higher in white as compared to gray matter regions (Figure 9C), NG2^+^ cells in white matter are not morphologically more complex than gray matter NG2^+^ cells, indicating that interaction with NOR is not a major factor shaping NG2^+^ cell morphology. In addition, when examining the end points of NG2^+^ cell processes, only 3-9% (Figure 8C) of process tips were found in close proximity to NOR which does not support the hypothesis that these processes extend specifically to contact NOR. NG2^+^ cells in corpus callosum were found previously to exhibit a higher degree of synaptic connectivity than NG2^+^ cells in hippocampus, cerebellar molecular layer, or cerebellar white matter (De Biase et al., 2010), yet callosal NG2^+^ cells are not significantly larger or more complex than NG2^+^ cells in other brain regions, suggesting that formation of synapses is also not a major determinant of NG2^+^ cell morphology. Further study will be needed to elucidate what governs the extension and branching of NG2^+^ cell processes.

Our observation of sparse interaction between NG2^+^ cells and NOR is in disagreement with recent estimates that NG2^+^ cells contact 71 ± 8% of NOR in the rat anterior medullary velum (Hamilton et al., 2010) and 33-49% of NOR in mouse corpus callosum, mouse spinal cord, and rat optic nerve (Serwanski et al., 2017). These differences may stem from distinct methodological approaches such as definitions of what constitutes “interaction” between NG2^+^ cells and NOR and from the inclusion of estimates of chance NOR-NG2+ cell interactions in our study. However, a more intriguing possibility is that these differences arise from analysis of different developmental stages. While we analyzed adult tissue (P55 – 85) in which myelination should be complete, Hamilton et al. and Serwanski et al. examined earlier developmental time points (P15 and P30, respectively). In addition, ankyrinG, which Hamilton et al. and Serwanski et al. used to identify NOR for quantification, has been shown to cluster at developing NOR prior to clustering of NaV and KV, and formation of a fully mature node (Dzhashiashvili et al., 2007). If NG2^+^ cells play a role in the early formation of immature NOR, this could explain the observation of a higher percentage of interactions at earlier developmental time points. Further, the small number of lingering interactions observed in the adult CNS could be due to ongoing myelin replacement and internode shortening that occurs throughout life. To explore this possibility we performed a limited analysis of optic nerve tissue from P25 mice and found that the percentage of NOR that come in close proximity to NG2^+^ cell processes did not differ significantly from adult mice (P = 0.17 Adult vs. P25 ON, corrected % NOR within 0.5 μm; P = 0.09 Adult vs. P25 ON, corrected % NOR within 1 μm; t-test)(Figure 10), suggesting that even before myelination of optic nerve is complete, only sparse interaction between NG2^+^ cells and NOR is observed.

**Figure 10.**
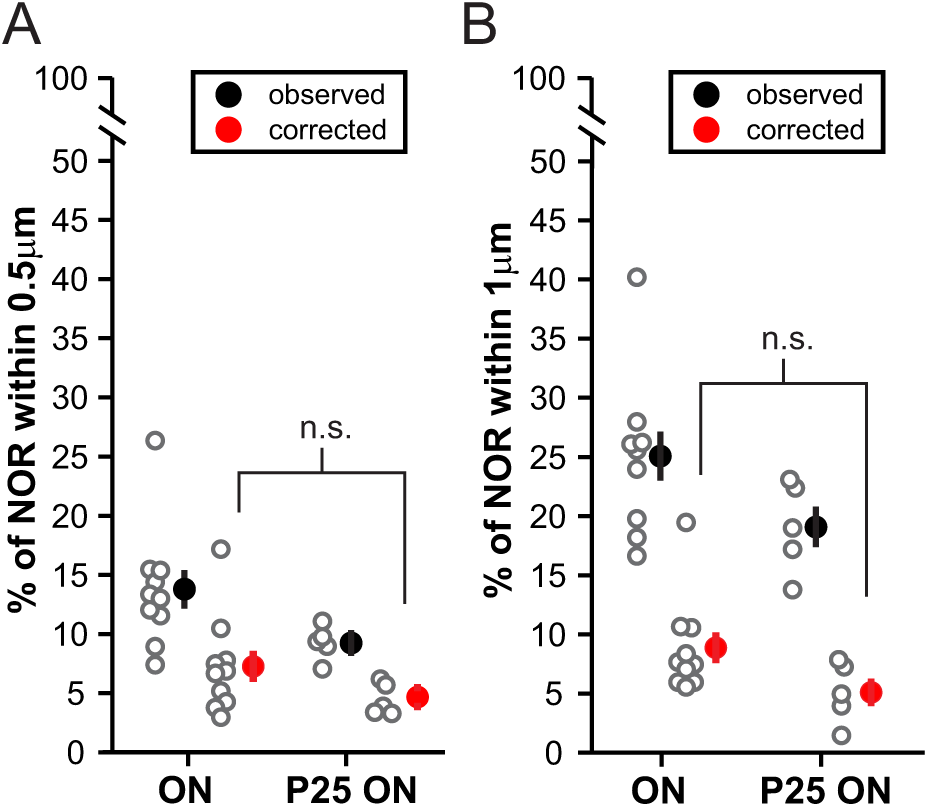
Spatial relationship between NG2+ cells and NOR in optic nerve from juvenile mice. ***A***, Percentage of NOR that come within 0.5 μm of the processes of optic nerve (ON) NG2^+^ cells in adult and juvenile mice. *Red circles* show the number of spatial interactions after correcting for interactions expected due to chance; P = 0.17 Adult vs. P25 ON, corrected % NOR within 0.5 μm, t-test. ***B***, Percentage of NOR that come within 1.0 μm of NG2^+^ cell processes in adult and juvenile mice. *Red circles* show the number of spatial interactions after correcting for interactions expected due to chance; P = 0.09 Adult vs. P25 ON, corrected % NOR within 1 μm; t-test.

What alternative hypotheses could explain the observation of sparse interaction between NG2^+^ cells and NOR? As oligodendrocyte progenitors, it could be that NG2^+^ cells need to sample activity from a certain, random subset of axons in their vicinity, in order to know whether additional myelination is necessary. Alternatively, perhaps neighboring NG2^+^ cells compete for a myelination-inducing signal and, as one neighbor wins out and begins to myelinate, the contact between the losing NG2^+^ progenitor and the NOR represents the remnant of this competition. It should also be pointed out that even in a homogeneous white matter tract such as the optic nerve, axons arise from distinct types of retinal ganglion cells that display different rates of activity. Perhaps NG2^+^ cells preferentially interact with the NOR of axons displaying certain activity patterns; findings from Serwanski et al. suggest that NG2 immunostaining is more frequently observed at NOR on large diameter axons and at NOR with larger nodal gap width. Finally, a caveat of our study, and all existing examinations of NG2^+^ cell-NOR interactions, is that these data represent observations at discrete moments in time. Using *in vivo* 2 photon imaging approaches, NG2^+^ cells in the cortex have been shown to actively remodel their processes over the course of minutes (Hughes et al., 2013). The present study cannot rule out the possibility that NG2^+^ cells dynamically contact NOR, to sample activity or deposit some signaling factor. However, the existence of NG2^+^ cell process “voids” which can span up to 25 μm suggests that large scale process remodeling would be necessary to contact some nodes. Fiber optic or GRIN lens technology, which will allow *in vivo* two photon imaging in deeper white matter structures, in combination with genetic tools such as the *NG2-mEGFP* mice will be necessary to address this possibility. In addition, genetic strategies aimed at ablation of NG2^+^ cells will provide the opportunity to clarify whether these cells play a role in regulating collateralization at NOR or in influencing axon conduction.

## Acknowledgements

The authors would like to thank: Dr. Jennifer Ziskin, Grace Kim, and Nadia Fakhari for help with pilot experiments and early development of experimental and analysis strategy; Dr. Martin Paukert for assistance with modifying MatLab code within Imaris MatLab XTensions; Dr. Richard Huganir (Johns Hopkins), Dr. Masahiro Fukaya (Hokkaido University), and Dr. Manzoor Bhat (UNC) for generously providing antibodies; The Multiphoton Imaging/Electrophysiology Core in the Department of Neuroscience at Johns Hopkins School of Medicine

## Notes

**Conflicts of Interest:** The authors report no conflict of interest.

